# Amyloid beta is released by lysosomal exocytosis from hiPSC-derived neurons

**DOI:** 10.1101/2025.03.01.640950

**Authors:** Adrianna R Tsang, Manoj Reddy Medapati, Claudia Seah, Jordan M Krupa, Stephen H. Pasternak

**Author notes:** Corresponding Author, RRI 3243, Robarts Research Institute, Schulich School of Medicine & Dentistry, Western University, 1151 Richmond St. N. London, ON N6A 5B7.

## Abstract

The endosomal-lysosomal system has long been linked to the production of amyloid beta (Aβ), but the specific intracellular compartments involved in Aβ secretion remain contentious. While lysosomes are typically associated with Aβ degradation, studies have also shown that lysosomes also serve as site of Aβ accumulation in cells, mouse models, and human tissues. Lysosomal exocytosis is a major secretory pathway in non-neuronal cells, but few studies have investigated this pathway in neurons. Here, we examined the potential role and mechanism of lysosomal exocytosis in human induced pluripotent stem cell (hiPSC)-derived neurons, and we hypothesized that lysosomal exocytosis is a pathway for Aβ secretion from these neurons. Using total internal reflection fluorescence (TIRF) microscopy, lysosomes filled with fluorescently labelled amyloid were seen approaching and fusing with the plasma membrane in real-time. The number and composition of the released particles were characterized using nanoscale flow cytometry. Silencing two proteins, Rab27b and munc13-4, significantly reduced these events and blocked the release of amyloid into the extracellular space. Our results provide direct evidence for the involvement of lysosomal exocytosis in the release of Aβ from neurons and highlight its potential as a target for therapeutic intervention in Alzheimer’s disease.

## Introduction

Amyloid deposition is one of the pathological hallmarks of Alzheimer’s disease (AD), the most common age-related neurodegenerative disorder worldwide. Although the exact cause of AD pathology is unknown, it is widely accepted that the accumulation of aggregated amyloid beta (Aβ) and tau-containing neurofibrillary tangles are central to AD. Amyloidogenic processing of Aβ by sequential cleavages from β-secretase BACE1 and ɣ-secretase leads to the generation of toxic Aβ peptides, Aβ_40_ and Aβ_42_.^1^ The amyloidogenic processing of the amyloid precursor protein (APP) is believed to occur after APP has been internalized into the endosomal-lysosomal system. Our group has shown that APP can be transported directly to lysosomes and processed into Aβ.^2–4^

Lysosomes are acidic compartments that are typically viewed as end state organelles responsible for degradation.^5^ Lysosomal dysfunction and the accumulation of intralysosomal Aβ have been implicated in progression of AD.^6^ Perhaps the earliest connection between lysosomes and AD was over 60 years ago when high levels of phosphatase-positive lysosomal dense bodies were localized within amyloid plaques of AD brains.^7^ Early electron microscopy studies also revealed swollen neurites filled with lysosome-like organelles surrounding Aβ plaques in the brains of AD patients.^8^ These initial findings have been supported by identification of enlarged or swollen lysosomes in neurons from post-mortem AD brain samples and mouse models of AD.^9–12^

In addition to digesting and recycling material, lysosomes are dynamic compartments that can alter their position in the cell to sense and respond to cellular needs in order to maintain homeostasis.^13,14^ Lysosomes also serve as secretory organelles undergoing Ca^2+^ dependent exocytosis in many cell types.^13,15–18^ For example, lysosomes function as major secretory granules in many haematopoietic cells, as well as osteoclasts and melanocytes.^16,19–26^ Cytotoxic granules in T-cells, basophil cell granules in basophils, azurophil granules in neutrophiles, and platelet dense in platelets are also key examples of cell-specific secretory lysosomes. The presence of secretory lysosomes in neurons and astrocytes has also been reported.^27–33^ As neuronal exocytosis of lysosomes is still an emerging field, most of our knowledge of lysosomal exocytosis has been studied in non-neuronal cells, including cargo, function and machinery. Exocytosis of lysosomes requires the recruitment of specific proteins which mediate their movement, tethering, and fusion to the plasma membrane.

Lysosomal exocytosis typically begins with a rise in intracellular Ca^2+^ that is detected by a Ca^2+^ sensor, synaptotagmin VII (SytVII).^20,21,27,34^ This triggers the mobilization of the lysosome toward the cell surface via a series of kinesin, actin and myosin proteins^27,35–37^ At the cell surface, the lysosome tethers and fuses with the plasma membrane through specific docking and fusion machinery, such as Soluble *N*-Ethylmaleimide-Sensitive Factor Attachment Receptor (SNARE) complexes and small GTPases of the Ras analog in brain (Rab) family.^15,16,26,27,35,38,39^ The fusion of lysosomal and plasma membranes results in the release of the original luminal cargo into the extracellular space. Many proteins have been implicated in this fusion, including v-SNARE, VAMP7, t-SNAREs, SNAP23 and syntaxin-4, and the small GTPases, Rab3 and Rab27.^15,23,35,39^ While none of these are unique to lysosomal exocytosis, Rab3 and Rab27 are the two so-called “secretory Rabs” and are present in neurons.^40–43^ Of these, Rab27 has been shown to be more stably localized at vesicle membranes.^44,45^ Rab27 exists as two isoforms, Rab27a and Rab27b, in which Rab27b displays higher localized expression in the central nervous system (CNS).^39,46^ In neurons, Rab27b targets and recruits its effector protein unc-13 homolog D (munc13-4), which plays a key role in priming of exocytic vesicles by interacting with SNARE complexes.^43^

As lysosomal dysfunction is linked to AD, the role of lysosomal exocytosis in the context of AD is an important field of research that is relatively understudied. It has been proposed that lysosomal exocytosis may serve as a prion-like transmission mechanism in which Aβ, along with other toxic proteins, can spread between cells.^29,33,47^ A study by Annunziata et al. (2013) investigated the link between lysosomal exocytosis and AD using knockout mice of neuraminidase 1 (NEU1), which displayed higher levels of lysosomal exocytosis, leading to an increase in Aβ secretion.^29^ Besides this study, not much is known about the regulated secretion of Aβ from neurons and this secretory event has never been directly imaged. Here, using human induced pluripotent stem cell (hiPSC)-derived neurons, we sought to visualize the real time release of Aβ by lysosomal exocytosis and investigate key machinery involved in this process, specifically focusing on Rab27b and its effector munc13-4.

## Results

### Aβ is preferentially localized in neuronal lysosomes

iPSC-derived human neurons were generated and differentiated using standard techniques adapted from Stem Cell Technologies (Supplementary Figure 1). To determine the intracellular localization of Aβ, wild-type iNeurons were loaded with exogenous fluorescently labelled Aβ_42_ monomers or APP-Swe iNeurons were incubated with Amytracker-520 to stain endogenous Aβ. The localization of Aβ was examined by the colocalization with transduced fluorescently-tagged compartment markers including, Rab5 for early endosomes, Rab9 for late endosomes, and LAMP1 for lysosomes (Figure 1). In wild-type iNeurons, only 15.61% of exogenously loaded Aβ_42_ co-localized with Rab5 positive compartments. By comparison, 74.78% of Aβ_42_ co-localized with Rab9 positive compartments (p=0.001) and 87.00% of Aβ_42_ co-localized with LAMP1 positive compartments (p<0.0001). However, no significant difference was seen between Rab9 late endosomes and LAMP1 lysosomes (p=0.38). On the other hand, in APP-Swe iNeurons, 87.92% of intracellular Aβ marked by Amytracker-520 co-localized with LAMP1 positive compartments. This was significantly higher than either endosomal marker, with only 5.30% Amytracker-520 co-localizing with Rab5 positive compartments (p<0.001) and 20.16% with Rab9 positive compartments (p<0.001).

**Figure 1.**
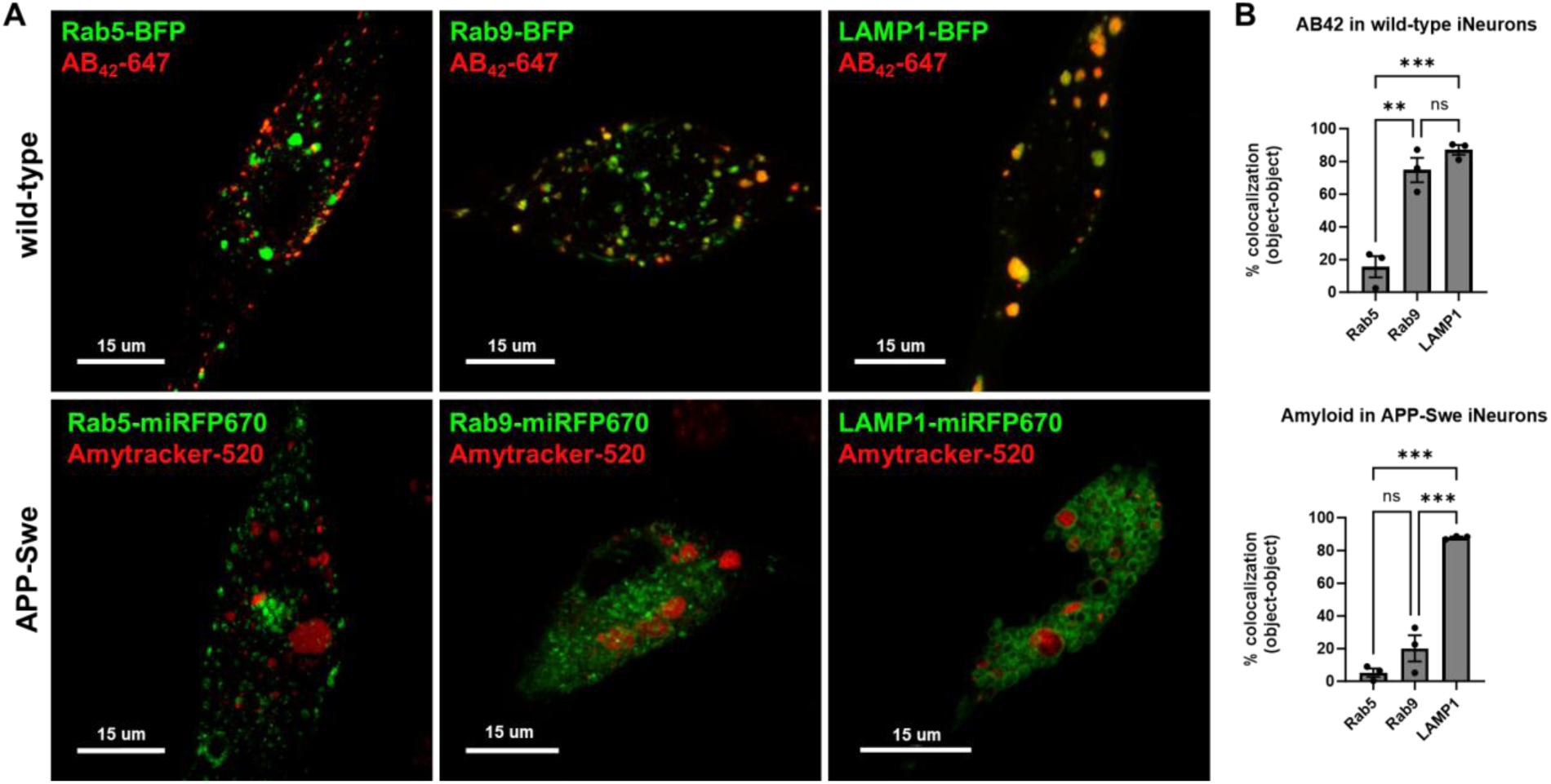
Live cell imaging of amyloid with compartment markers in wild-type and APP-Swe iNeurons. A) Representative images of wild-type cells (top) loaded with Aβ_42_-647 (red) or APP-Swe cells (bottom) incubated with Amytracker-520 (red). Cells were transduced with adenoviral vectors containing fluorescently tagged Rab5, Rab9, LAMP1 to mark early endosomes, late endosomes and lysosomes, respectively (green). B) Object-object based colocalization was calculated using the spot and surfaces functions on Imaris. All results are presented as mean ± SEM. Asterisks indicate significant changes according to one-way ANOVA and post hoc Tukey test. **p<0.01 ***p<0.001

This supports the idea that accumulation of aggregated Aβ occurs within lysosomes in AD. The non-significant differences between late endosomes and lysosomes, may be due to the overlapping expression of Rab9 and LAMP1. Thus, dextran loading was used to more accurately identify terminal lysosomes.^48^ Wild-type iNeurons were incubated with 10 kDa dextran-TMR for 6 hours, followed by an overnight incubation at 37℃ for 20 hours (Figure 2). The timing of this pulse-chase was used to ensure that the dextran had fully traversed the endocytic pathway and was confined to lysosomes, which has been previously demonstrated.^49^ Across all compartment markers, 88.19% of all Aβ_42_ spots contained dextran and 92.39% contained LAMP-1. Importantly, when Rab5 and Rab9 compartment markers were exclusively separated from dextran signal (i.e, Rab5(+) dex(-) and Rab9(+) and dex(-)), the percentage of Aβ_42_ spots significantly fell to 3.18% and 4.93% for Rab5 (p<0.001) and Rab9 (p<0.001), respectively. This strongly supports the idea that Aβ_42_ accumulates primarily in lysosomes in AD.

**Figure 2.**
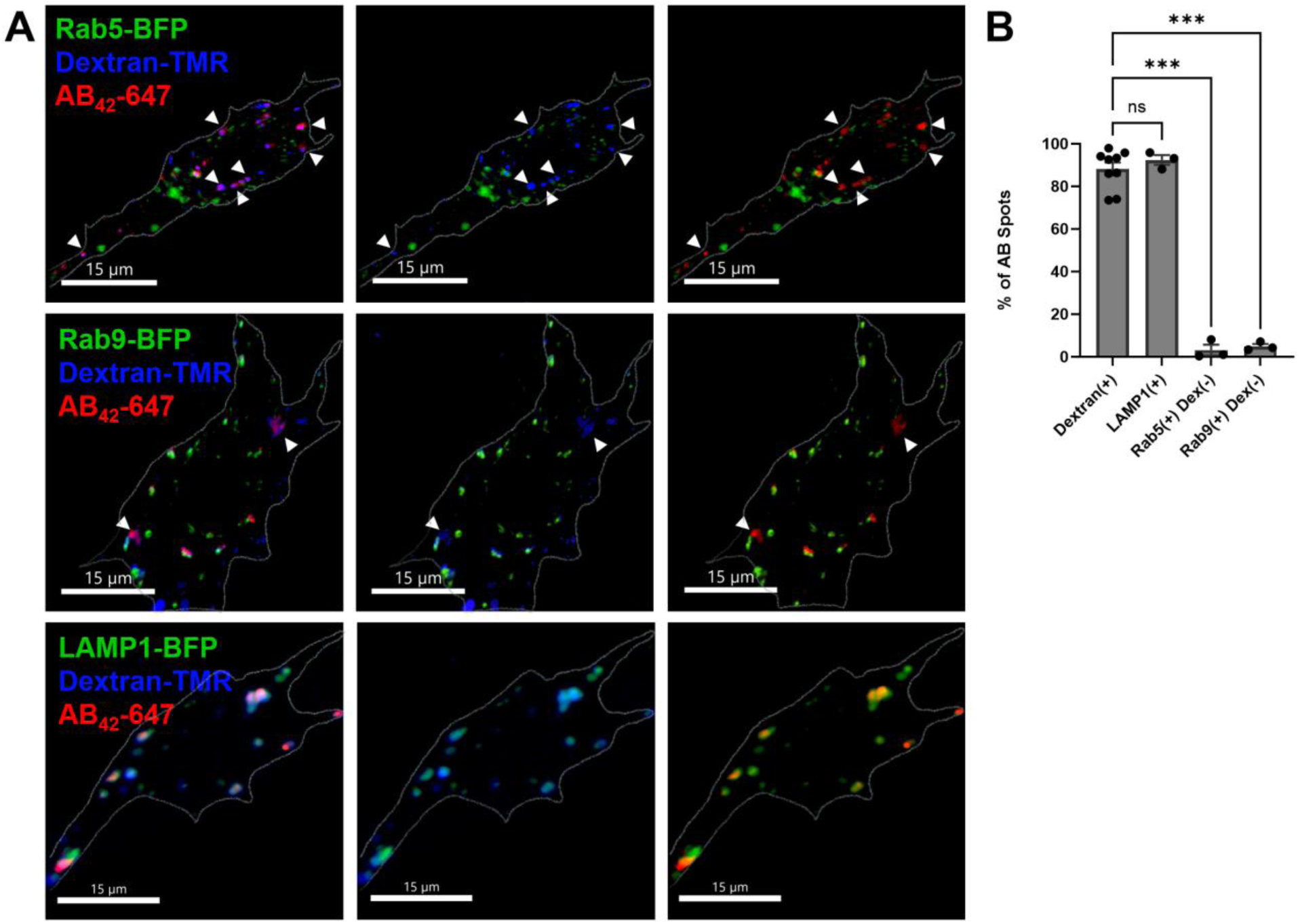
Live cell imaging of amyloid with dextran in wild-type iNeurons. A) Representative images of wild-type cells loaded with Aβ_42_-647 (red). Cells were transduced with adenoviral vectors containing fluorescently tagged Rab5, Rab9, LAMP1 to mark early endosomes, late endosomes and lysosomes, respectively (green). A 6-hour pulse with dextran and 20 hour chase was used to ensure that the probe was confined to lysosomes (blue). Arrowheads represent Aβ_42_ with dextran positive signal but lacking Rab5 or Rab9 signal. B) Classification and quantification of Aβ_42_ spots containing dextran and/or compartment markers signals was calculated by Imaris. All results are presented as mean ± SEM. Asterisks indicate significant changes according to one-way ANOVA and post hoc Tukey test. ***p<0.001

### TIRF microscopy captures lysosomal release of amyloid

To follow lysosomal trafficking, iNeurons were transduced with LAMP1-BFP to label lysosomes and pHmScarlet-LAMP1 to monitor intraluminal pH. pHmScarlet exhibits minimal fluorescence under acidic conditions but demonstrates strong fluorescence at neutral pH. Upon fusion of lysosomes with the plasma membrane, the incoming culture media neutralizes the acidic environment of the lysosomes. This shift to a neutral pH triggers a marked increase in fluorescence of pHmScarlet.

The changes in fluorescence intensity of individual lysosomes were captured using total internal reflection fluorescence (TIRF) microscopy. In brief, TIRF microscopy provides higher resolution of events occurring near the plasma membrane by restricting the depth of light energy able to penetrate a cell. To do this, laser light is reflected off the surface of the glass coverslip at a critical reflection angle in which almost all of the energy is reflected. However, an evanescent wave that can excite fluorophores penetrates 70-250 nm into the specimen surface. This eliminates any background signal outside this specific z-plane that would be captured under normal epifluorescence microscopy and allows for high-resolution time-lapse videos.

At rest, exogenously loaded fluorescent Aβ_42_ and endogenous Amytracker-stained Aβ_42_ was seen in LAMP1 compartments that had little pHmScarlet signal, indicating that they were acidified lysosomes captured by live-cell TIRF microscopy (Figure 3). Ionomycin, a Ca^2+^ ionophore, was used to trigger exocytosis, and live-cell videos were acquired in wild-type cells (Supplementary Movie 1) and APP-Swe cells (Supplementary Movie 2).^21,28,35,50^ In wild-type iNeurons loaded with Aβ_42_, lysosomes began fusing with the plasma membrane 10-20 seconds following addition of ionomycin, resulting in a gain in pHmScarlet-LAMP1 intensity. Meanwhile, there was a simultaneous reduction in Aβ_42_ intensity within LAMP1-BFP positive compartments. In APP-Swe cells labelled with Amytracker-520, there was an increase in pHmScarlet-LAMP1 intensity for approximately 30 seconds following ionomycin treatment, along with a simultaneous decrease of LAMP1-BFP and Amytracker-520 intensities.

**Figure 3.**
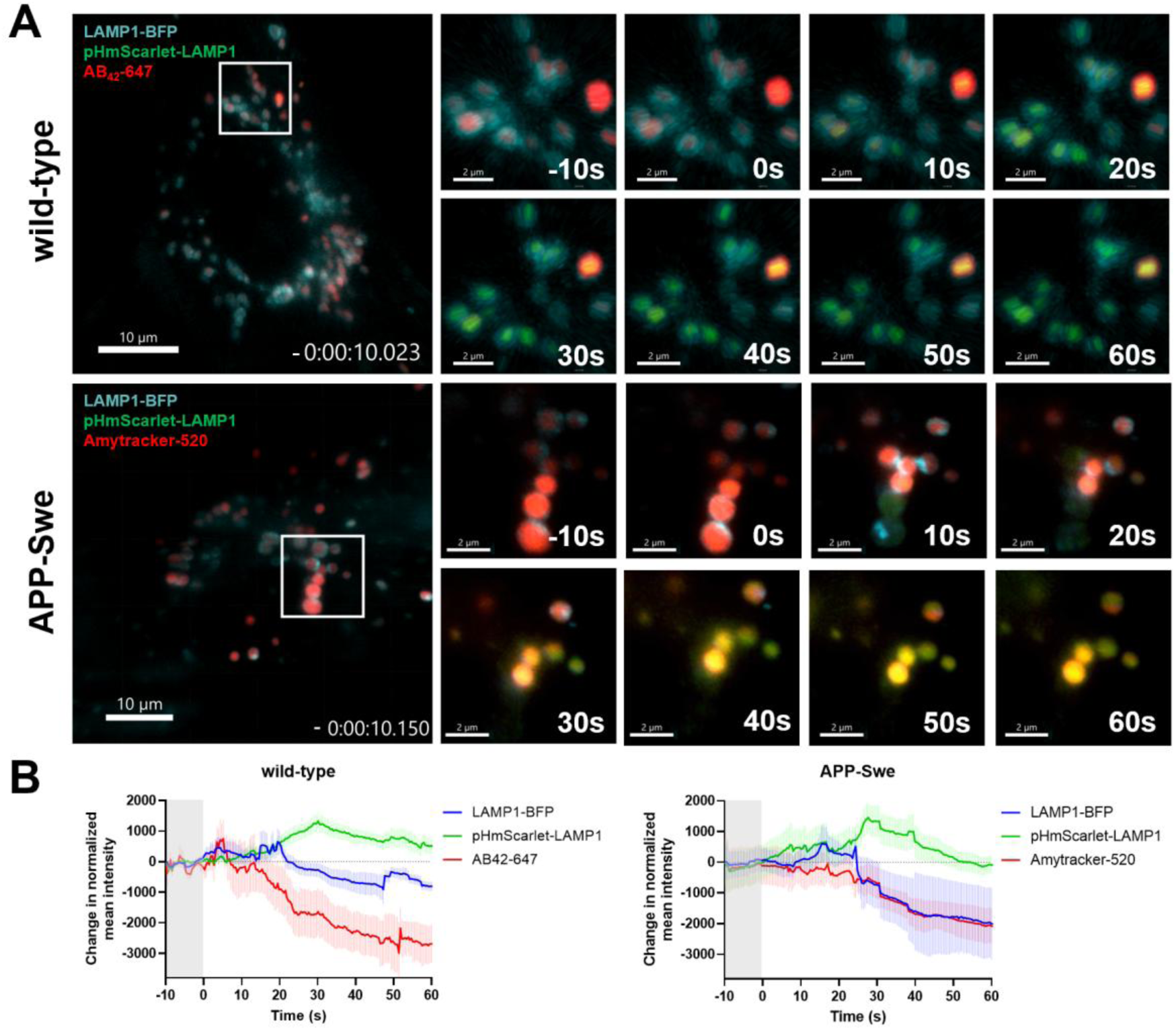
TIRF microscopy of amyloid released from lysosomes in wild-type and APP-Swe iNeurons. A) Representative images from a live cell video captured through a TIRF microscope demonstrating lysosomal exocytosis in a wild-type and APP-Swe cell after ionomycin treatment over 60 seconds. Cells were transduced with adenoviral vectors containing fluorescently tagged LAMP1-BFP (blue) and pHmScarlet-LAMP1 (green) and incubated with Aβ (red). B) Change in normalized mean intensity of each channel was tracked after ionomycin treatment (at time = 0s). A total of 538 lysosomes were analyzed in wild-type cells (n=10), and 785 lysosomes were tracked in APP-Swe cells (n=10).

### Silencing of Rab27b and munc13-4 impedes lysosomal exocytosis of amyloid

To study the machinery involved in lysosomal exocytosis in neurons, silencing of Rab27b and munc13-4 was performed using shRNA adenoviruses (Supplementary Figure 2). Silencing of both proteins in wild-type and APP-Swe iNeurons completely abolished the rise in the pHmScarlet signal captured by TIRF microscopy (Figure 4). The intensities of LAMP1-BFP, pHmScarlet-LAMP1, Aβ_42_-647 or Amytracker-520 all remained relatively stable. This lack of intensity changes represented a failure of LAMP1-positive vesicles to fuse with the plasma membrane and thus, prevent the release of Aβ into the extracellular space.

**Figure 4.**
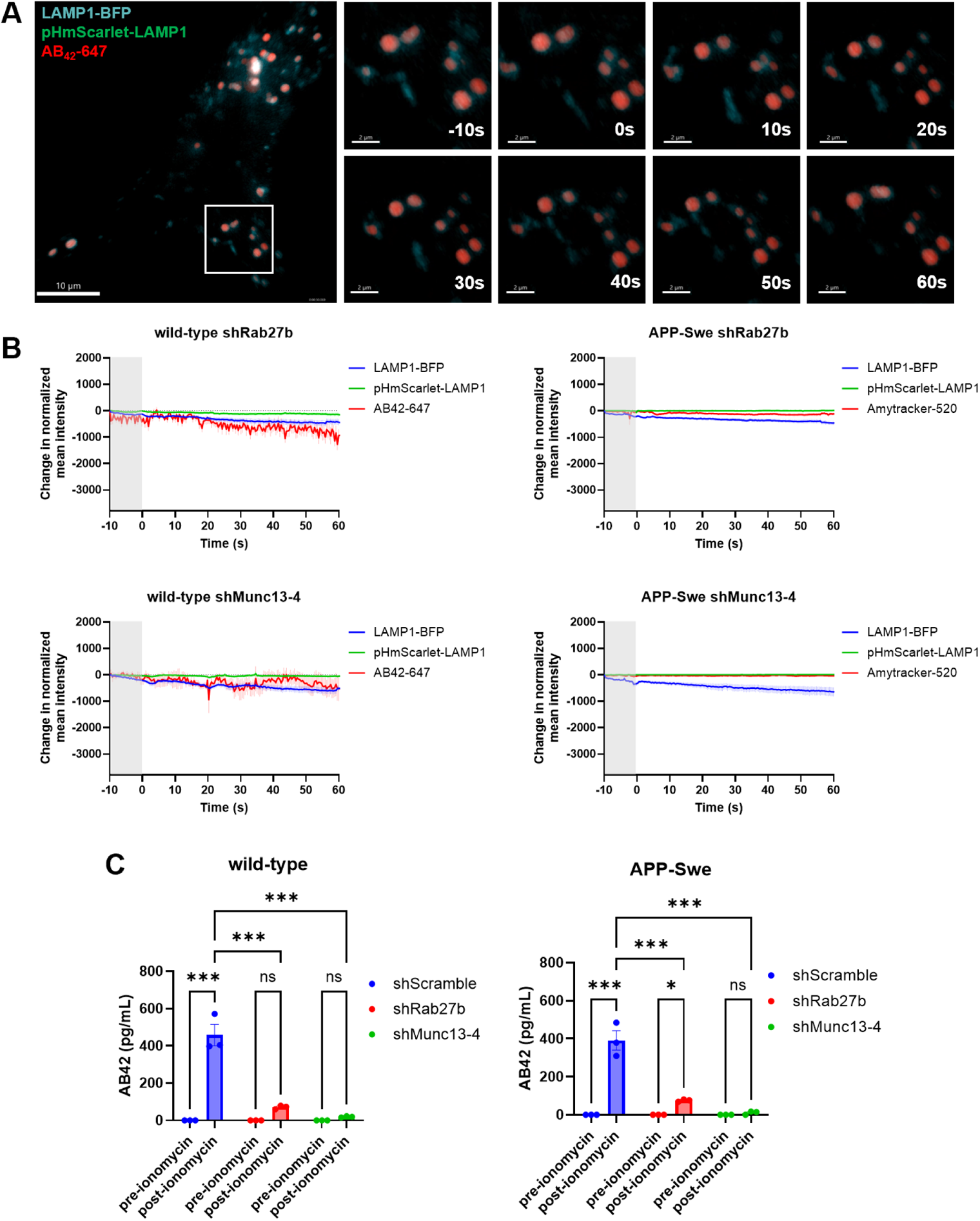
Silencing of Rab27b and munc13-4 halts lysosomal exocytosis. A) Representative images from a live cell video captured through a TIRF microscope demonstrating the lack of lysosomal exocytosis in a wild-type cell upon shRab27b silencing. B) Silencing of Rab27b and munc-13-4 results in intensities across all 3 channels to remain relatively stable, demonstrating a lack of lysosomal secretion. C) Significant reduction of Aβ is detected in the media of iNeurons following silencing, as measured by ELISAs. All results are presented as mean ± SEM. Asterisks indicate significant changes according to two-way ANOVA with post hoc Tukey test. *p<0.05 ***p<0.001

To verify that Aβ was released during ionomycin-induced exocytosis and that this release was blocked by silencing of Rab27b and munc13-4, media samples were collected before and after exocytosis, and Aβ levels were analyzed by ELISAs. Wild-type and APP-Swe cells demonstrated a significant increase in Aβ release following ionomycin treatment. In wild-type cells, the levels of extracellular Aβ_42_ rose from undetectable levels to 458.33 pg/mL following ionomycin treatment (p<0.001). This was reduced to 70.77 pg/mL by silencing of Rab27b (p<0.001) and to 19.67 by silencing of munc13-4 (p<0.001). Likewise, this was also seen in APP-Swe iNeurons, in levels of extracellular Aβ_42_ rose from undetectable levels to 391 pg/mL following ionomycin treatment (p<0.001). This was reduced to 75.33 pg/mL from shRab27b treated cells (p<0.001) and to 10.67 pg/mL from shMunc13-4 treated cells (p<0.001).

### Lysosomes are the main secretory organelle releasing amyloid-containing particles upon ionomycin treatment

To better examine the particles that are released in the extracellular space, nanoscale flow cytometry (or nanoflow analysis) was performed using an Apogee A60 Microplus system, which is optimized for the detection of 80-100 nm range particles with much higher sensitivity. We have previously used this system to identify tau and amyloid-containing particles from AD patient plasma.^51^ iNeurons were transduced with fluorescently-tagged compartment markers and either loaded with fluorescent Aβ (for wild-type iNeurons) or labelled with Amytracker (for APP-Swe iNeurons) (Figure 5).

**Figure 5.**
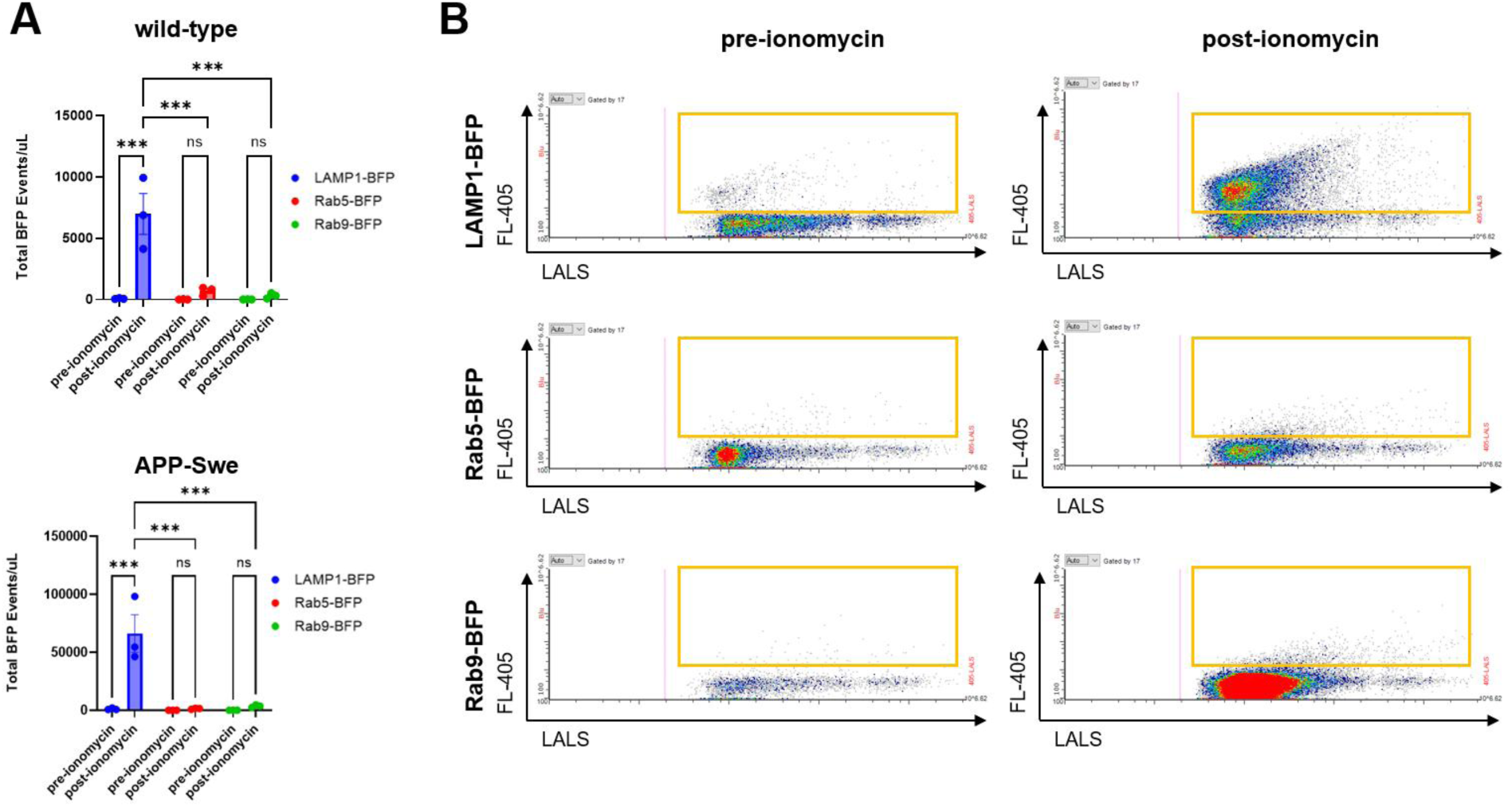
Nanoscale flow cytometry of secreted vesicles triggered by ionomycin. A) Total BFP events/µL were analyzed by nanoscale flow cytometry on media of wild-type and APP-Swe iNeurons following ionomycin treatment. B) Representative cytograms of LAMP1-BFP, Rab5-BFP and Rab9-BFP collected from cell culture media prior and following ionomycin treatment. All results are presented as mean ± SEM. Asterisks indicate significant changes according to two-way ANOVA and post hoc Tukey test. ***p<0.001

The media was collected pre- and post-ionomycin treatment and fluorescent particles in the media were sorted by nanoscale flow cytometry. Prior to ionomycin treatment, almost no fluorescent particles were detected in the media. Following ionomycin treatment to trigger exocytosis, majority of BFP-containing particles were detected from cells expressing LAMP1-BFP, but few from cells transduced with Rab5-BFP or Rab9-BFP. In wild-type cells, an average of 6981.89 events/µL LAMP1 containing particles was detected in samples post-ionomycin treatment, which is a significant increase from 77.33 events/µL pre-ionomycin treatment (p<0.001). In contrast, no significant levels of Rab5 (688.16 events/µL, p=0.5) or Rab9 (305.91 events/µL, p=0.77) were measured in the media following ionomycin treatment. In APP-Swe cells, an average of 66338 events/µL LAMP1 containing particles was measured in samples post-ionomycin treatment, which was a significant increase from 1138.22 events/µL pre-ionomycin treatment (p<0.001). No significant changes were detected for Rab5 (1416.33 events/µL, p=0.89) or Rab9 (3583 events/µL, p=0.72).

### Nanoscale flow cytometry reveals differences in Aβ-containing particles released from wild-type and APP-Swe iNeurons

Interestingly, our nanoscale flow cytometry results also found a stark difference between the type of Aβ-containing particles released by wild-type versus APP-Swedish iNeurons (Figure 6). In addition to detecting LAMP1, we also examined the presence of common exosomal marker CD9. In wild-type cells, 52.1% of smaller fluorescent Aβ-containing particles (180-300 nm) did not contain LAMP1 or CD9 (LAMP1-/CD9-). This was significantly greater than Aβ secreted which contained LAMP1 only (15.2%, p<0.001), CD9 only (3.73%, p<0.001), or with LAMP1 and CD9 together (21.2%, p<0.001). In contrast, 57.3% of larger Aβ-containing particles (300-1000 nm) was secreted with LAMP1 and CD9 together, which was significantly greater than Aβ secreted with LAMP1 only (10.7%, p<0.001), CD9 only (4.78%, p<0.001), or as unbound cargo (21.1%, p<0.001).

**Figure 6.**
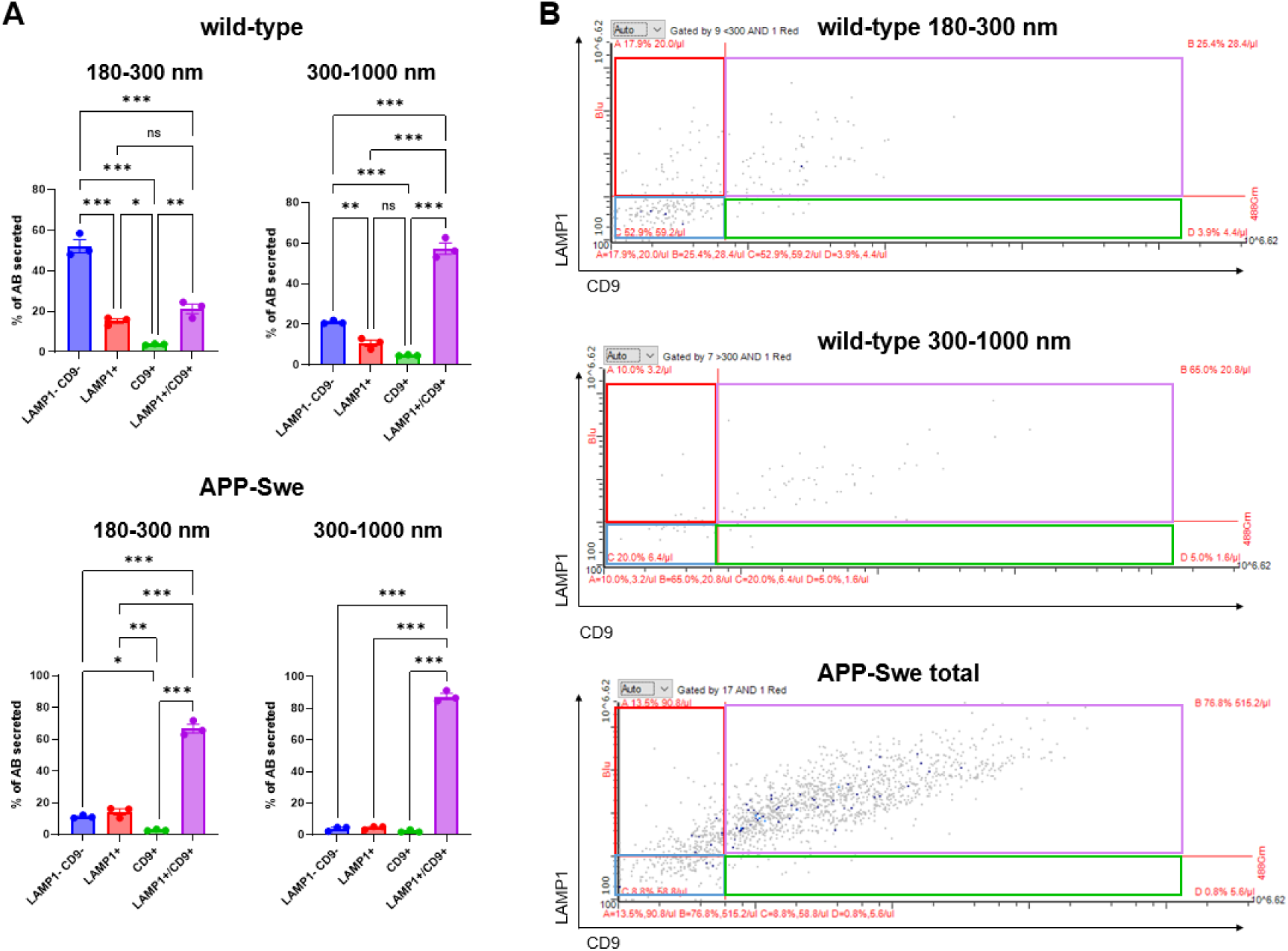
Nanoscale flow cytometry of particles containing Aβ. B) Percentage of unbound Aβ (LAMP1-/CD9-), Aβ with LAMP1, CD9 or both (LAMP1+/CD9+) was measured by nanoscale flow cytometry on media from wild-type and APP-Swe iNeurons following ionomycin treatment. Particles were sorted by size (established with calibration beads). B) Representative cytograms of Aβ collected from cell culture media following ionomycin treatment. Blue boxes indicate particles containing Aβ only, red boxes indicate particles of Aβ with LAMP1, green boxes indicate particles of Aβ with CD9, and purple boxes indicate particles of Aβ with LAMP1 and CD9. All results are presented as mean ± SEM. Asterisks indicate significant changes according to one-way ANOVA and post hoc Tukey test. *p<0.05, **p<0.01. ***p<0.001

In APP-Swedish cells, the majority of Amytracker-labelled Aβ was released in particles that contained LAMP1 and CD9 together, regardless of size. 66.7% of smaller Aβ (180-300 nm) was secreted in particles bearing LAMP1 and CD9, which was significantly greater than Aβ secreted with LAMP1 only (14.2%, p<0.001), CD9 only (2.78%, p<0.001), or as unbound cargo (11.2%, p<0.001). Likewise, 87.5% of larger Aβ (300-1000 nm) was secreted in particles containing both LAMP1 and CD9, which was significantly greater than Aβ secreted bearing LAMP1 alone (4.49%, p<0.001), CD9 alone (2.27%, p<0.001), or lacking both (3.89%, p<0.001).

These findings suggest that small exogenously loaded Aβ_42_ aggregates are released as unbound cargo in wild-type cells, whereas aggregated Aβ is mainly released within membrane bound compartments in APP-Swe cells.

## Discussion

Despite extensive research on Aβ, direct imaging of Aβ secretion in human neurons has never been shown. Prior work in our lab has established the involvement of endosomes and lysosomes in Aβ production.^2–4,52^ Here, we confirmed that exogenously loaded fluorescent-labelled Aβ and endogenously produced Aβ resides within lysosomes. Using TIRF microscopy, we directly observed exogenously-loaded fluorescent Aβ_42_ release via lysosomal exocytosis following ionomycin treatment. Similarly, the secretion of endogenous Aβ, identified with Amytacker in APP-Swe cells, was also imaged by TIRF microscopy as it was secreted from the cells upon ionomycin induction. This was blocked by shRNA knockdown of genes known to participate in lysosomal secretion (Rab27b and Munc13-4), and confirmed by ELISA-based detection of Aβ in the media following stimulation. Nanoscale flow cytometry revealed that small Aβ-containing aggregates released in this process are predominantly released without the membrane/exosome markers LAMP1 or CD9, while larger aggregated Aβ is released within LAMP1/CD9 containing structures. The findings of this study demonstrated the exocytic capabilities of neuronal lysosomes, and that lysosomes release of Aβ and Aβ-containing aggregates into the extracellular space.

This adds to the limited research on the presence of secretory lysosomes in cells of the central nervous system and in neurons.^27–33^ However, most of our understanding of the machinery involved in this process comes from studies in non-neuronal cells.^16,19–26^ The involvement of Rab27b in lysosomal secretion is well-supported by research studies that underscore its critical role in vesicular trafficking and exocytosis.^42–45^ For instance, silencing of Rab27b results in the perinuclear accumulation of CD63, a marker for late endosomes and lysosomes.^39^ In the context of elimination of pathological substrates, Rab27b has been implicated in the exocytosis of alpha-synuclein.^53–55^ In particular, knocking down Rab27b disrupts lysosomal function, leading to the accumulation of defects in primary neuronal cultures and in the cortex and hippocampus of Rab27b knockout mice.^53^ Future work will be needed to map out the regulatory structure of lysosomal exocytosis in neurons, including the role of calcium sensors such as SytVII,^27,34^ and proteins of the SNARE machinery, particularly the t-SNARE syntaxin-4 and the v-SNARE VAMP-7 that are critical for the fusion of lysosomes with the plasma membrane.^24,35,56^ While Rab27b is a known secretory Rab that has been shown to be highly abundant in the central nervous system, only a handful of papers have examined Rab27 in AD.^40–43^

This work supports also the longstanding idea that lysosomes are important in the aggregation and release of Aβ. Our own work suggests that APP can be rapidly transported to lysosomes and from the plasma membrane by macropinocytosis, and directly from the Golgi apparatus, where it can be cleaved into Aβ.^2–4^ Even in the 1990s, there was evidence that extracellular Aβ is taken up into lysosomes, where it can nucleate endogenous Aβ aggregation.^57,58^ Aβ is known to accumulate and aggregate intracellularly in lysosomes.^11,12,59–61^ The aggregation of Aβ in late endosomes and lysosomes can disrupt synapses^62^ and cause lysosomal rupture leading to cell death.^63^ Aβ secretion is known to be driven by neural activity.^64^ Because lysosomal enzymes are only activated in the lysosomes, presence of active lysosomal enzymes in amyloid plaques suggests that they are derived from lysosomes.^65^

In summary, here we demonstrate that Aβ is directly secreted from neurons by lysosomal secretion. This work dovetails with longstanding observations of Aβ aggregation and accumulation in lysosomes and the observation that extracellular Aβ plaques contain active lysosomal enzymes. The secretion of Aβ in aggregates from lysosomal secretion suggests that this process may play a role in generating aggregates that could propagate in the development of AD.

## Methods

### Cell culturing of hiPSC-derived neurons

hiPSCs were obtained from the New York Stem Cell Foundation, including cell lines containing the APP Swedish mutation (APP-Swe; CO002-01-CS-003, Lot# E137-1D) and its isogenic control (wild-type; CO0002-01-SV-003, Lot# E259-1H). The induction of hiPSCs into neural progenitor cells (NPCs) was performed using STEMdiff™ SMADi neural induction kit (Stem Cell Technologies, 08581) and STEMdiff™ neural induction media (Stem Cell Technologies, 05835), following the manufacturer’s protocols outlined in ‘Generation and Culture of Neural Progenitor Cells Using the STEMdiff™ Neural System’. In brief, embryoid bodies were initially generated, maintained for 10 days, and replated for neural rosette selection using STEMdiff™ neural rosette selection reagent (Stem Cell Technologies, 05832). NPC outgrowths were then formed from selected rosette-containing clusters.

Selected NPCs were then maintained and passaged (every 5-7 days) in STEMdiff™ neural progenitor medium (Stem Cell Technologies, 05833), or differentiated for 7 days using STEMdiff™ forebrain neuron differentiation kit (Stem Cell Technologies, 08600). Following differentiation, cells were matured into iNeurons for 21 days in vitro (DIV) using Neurobasal™ medium (Gibco, 21103049) supplemented with 1x B27 (Gibco, 17504044), 0.5× GlutaMAX (Gibco, 35050061), 1x antibiotic-antimycotic (Gibco, 15240096), 0.5 mM db-cAMP (Sigma-Aldrich, D0627), 0.2 ug/mL ascorbic acid (Sigma-Aldrich, A92902), 20 ng/mL BDNF (PeproTech, 450-02), and 20 ng/mL GDNF (PeproTech, 450-10). For all experiments, cells were maintained or plated on cultureware coated with 15 ug/mL poly-L-ornithine (Sigma-Aldrich, P4957) and 10 ug/mL laminin (Corning, 354232).

### Viral vectors

All constructs were designed, packaged into adenovirus serotype 5, and purchased from VectorBuilder. iNeurons were transduced at 18 DIV with the listed multiplicity of infections (MOIs) and performed following recommendations set by VectorBuilder.

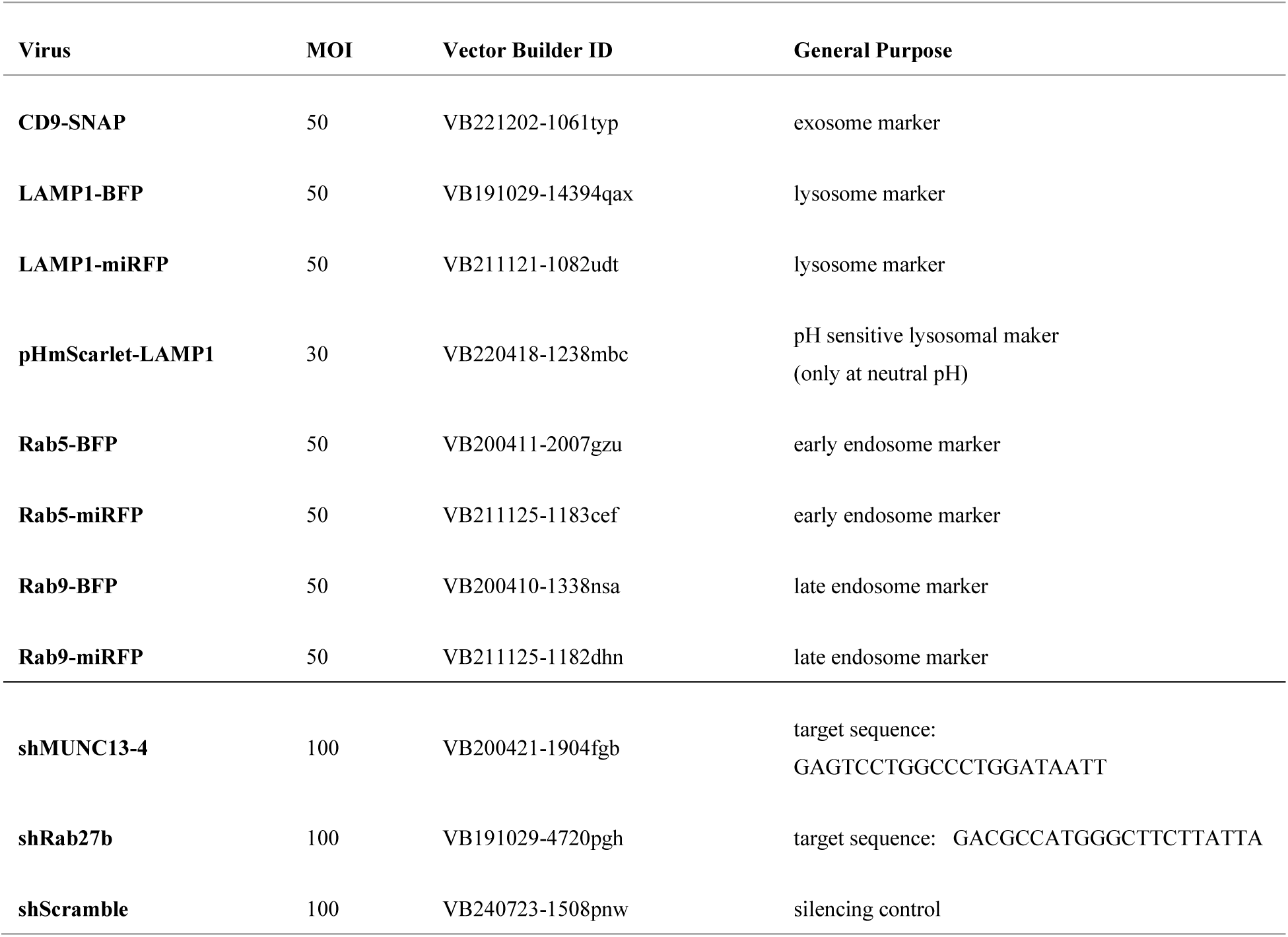

### Aβ monomer preparation and loading & Amytracker detection

Aβ_42_ monomers were generated from recombinant Aβ_42_ tagged with HiLyte™ Fluor 647 (Anaspec, AS-64161), following previously established protocols.^66^ In brief, lyophilized peptides were solubilized in 1,1,1,3,3,3-hexafluoro-2-propanol (Sigma-Aldrich, 105288). Vacuum centrifugation was used to evaporate the HFIP for subsequent resuspension in dimethyl sulfoxide (DMSO). Following a 10-minute period of sonication, Aβ_42_ monomers were diluted in ice-cold PBS. Matured wild-type iNeurons (20 DIV) were loaded with 250 nM of Aβ_42_ overnight prior to experimentation for total internal fluorescence (TIRF) microscopy.

Instead of Aβ_42_ loading, APP-Swe iNeurons were incubated with Amytracker-520 (1:250, Ebba Biotech) for 2 hours to mark endogenous aggregated Aβ for microscopy experiments. For nanoscale flow cytometry experiments, the media of APP-Swe iNeurons was incubated with Amytracker-680 (1:100, Ebba Biotech) for 10 minutes prior to analysis.

### Ionomycin treatment

Following Aβ_42_ loading overnight or Amytracker incubation for 2 hours, iNeurons were washed and treated with 5 or 10 µM of ionomycin (Sigma-Aldrich, C7522) in Hank’s Balanced Salt Solution (HBSS, Gibco, 14025092) to induce lysosomal exocytosis. 5 µM of ionomycin was used for microscopy experiments, whereas 10 µM of ionomycin was used for enzyme-linked immunosorbent assays (ELISAs) and nanoscale flow cytometry. This difference is mainly due to the fact that for microscopy experiments, ionomycin was directly injected over the confocal dish for live-cell capture of events, and thus a lower concentration was used, whereas HBSS containing ionomycin was replaced across the entire 6-well for ELISAs and nanoscale flow cytometry.

### Immunofluorescence staining

To verify the neural cell yield, immunofluorescence staining was performed for nestin and MAP2 (Supplementary Figure 1). In brief, iNeurons were fixed for 15 minutes with 4% paraformaldehyde. Cells were permeabilized for 10 minutes with 0.1% Triton X-100 in HBSS and blocked with 2% bovine serum albumin (BSA) for 30 mins. Cells were incubated with primary antibodies for MAP2 (1:1000; Sigma-Aldrich, M9942) and nestin (1:200; Invitrogen, PA5-11887) for 1 hour at RT, washed three times with HBSS, and incubated with species-appropriate secondary antibodies conjugated to Alexa Fluor 488 (1:1000; Invitrogen, A21202) or Alexa Fluor 647 (1:1000; Invitrogen, A31573) for 1 hour at RT. Cells were then counterstained with DAPI (Sigma-Aldrich, D9542).

### Confocal microscopy and image analysis

Images were acquired on a Leica TCS SP8 confocal microscope (Leica, Germany) using an HC PL APO CS2 63x/1.40 oil objective, with excitation lasers of 405 nm, 488 nm, 552 nm, and 638 nm. Emission filters were consistent for each set of experiments.

For immunofluorescence experiments with fixed cells, a minimum of 3 regions were acquired per confocal dish, with each region consisting of a stitched merge of 5×5 tiles. To quantify the neural yield of 3 experimental replicates, the relative percentage of nestin and MAP2 positive cells was calculated using the filament tracer function on Imaris 10 Software (Oxford Instruments).

For all co-localization experiments, Z-stack images of the entire cell were acquired from a minimum of 3 live-cells per confocal dish, per replicate. Imaris was used for semi-automated quantification across 3 experimental replicates. In Aβ and compartmental marker experiments, object-object based colocalization was measured by calculating the percentage of co-localization between spots (semi-automatically generated for Aβ) and surfaces (semi-automatically generated for compartment markers). In experiments using 10 kDa dextran-tetramethylrhodamine (TMR; Invitrogen, D1868), spots were created to represent Aβ_42_, and intensity filters were then used to classify spots based on the presence of dextran-TMR and/or compartment maker signals.

### TIRF microscopy

To capture live cell videos of individual lysosomes collapsing at the plasma membrane, total internal reflection fluorescence (TIRF) microscopy was performed using a Leica DMI 6000B TIRF microscope (Leica, Germany) with an HCX PL APO 63X oil immersion objective with numerical aperture of 1.47. Fluorescence emitted from excitation lasers at 405 nm, 488 nm, 561 nm, and 635 nm was captured after passing through a Quad-Band filter cube. The Quad-Band filter cube setup allowed for a very short switching time and thus, an ultra-high synchronized frame rate that is required for the analysis of TIRF videos.

Following Aβ_42_ loading overnight or Amytracker incubation for 2 hours, iNeurons were washed with pre-warmed HBSS. Cells were then transported to the microscope stage in the incubation chamber and treated directly with 5 µM of ionomycin. To capture rapid exocytic events, the imaging time for each wavelength was synchronized with the hardware. For every video, images were captured approximately every 0.3-0.5 seconds (across 3 channels) for 150-300 time points.

A minimum of 10 live cells were captured using TIRF microscopy and analyzed with Imaris 10 Software (Oxford Instruments). Using the semi-automated spot detection feature on Imaris, a spot was created to represent each lysosome (based on the pHmScarlet-LAMP1 channel), and tracks were generated to follow the fate of each lysosome over time. At each single time point, the mean intensity of each channel was measured from that spot and averaged across all spots within a single cell. Tracks were normalized to its maximum intensity, and the change in normalized intensity was plotted against time to compare across conditions.

### Western blotting

Western blotting for Rab27b and munc13-4 was performed to verify their silencing (Supplementary Figure 2). iNeurons were washed on ice with ice-cold HBSS and then lysed in N-PER lysis buffer (Thermo Scientific, 87792) with 1X protease and phosphatase inhibitor cocktail (Thermo Scientific, 78442). Protein concentrations were measured using the Pierce BCA Protein Assay (Thermo Scientific, 23225).

Equal amounts of total protein were mixed with 2X Laemmli buffer (Bio-Rad, 1610737), denatured at 95℃ for 5 minutes, and loaded onto 4-20% Mini-PROTEAN® TGX^™^ precast protein gels (Bio-Rad, 4561096). Gels were run at 125V for 75 minutes and subsequently, transferred onto methanol-activated Immun-Blot® PVDF membranes (Bio-Rad, 1620177) at 90 V for 75 minutes. Following Ponceau verification of transfer, membranes were blocked with 5% BSA for 1 hour. Membranes were incubated in primary antibodies (α-Rab27b: 1:500, Proteintech, 13412-1-AP, α-munc13-4: 1:500, Proteintech, 16905-1-AP, and α-alpha-tubulin: 1:100,000, Sigma-Aldrich, T5168) in blocking solution overnight at 4°C. Membranes were incubated with HRP-conjugated goat anti-rabbit (1:5000; Bio-Rad, 1706515) for 1 hour at RT, followed by visualization using Amersham ECL prime western blotting detection reagent (Cytvia, RPN2236) on a Bio-Rad ChemiDoc MP. Densitometric analysis was conducted using ImageLab Software (Bio-Rad).

### Enzyme-linked immunosorbent assays (ELISAs)

For ELISAs, matured iNeurons grown on coated 6-well plates were washed with pre-warmed HBSS and incubated with 10 µM ionomycin in HBSS at 37℃ for 5 minutes prior to supernatant collection. Samples were centrifuged at 500 x g for 5 minutes to remove any debris and the supernatant was then collected and frozen at - 80℃. For measurement of Aβ_42_ levels, supernatants were diluted 10-fold and detected using an ELISA for human Aβ_42_ (Invitrogen, KHB3544). Triplicates of each sample from 3 replicates were loaded per plate.

### Nanoscale flow cytometry

Supernatant samples containing exocytosed contents were collected similarly to the ELISA samples, as described above. For detection of CD9-SNAP in supernatants by nanoscale flow cytometry, labelling was performed using SNAP-Cell™ Oregon-Green (New England Biolabs, S9104S) according to the manufacturer’s instructions.

Undiluted samples were analyzed on the Apogee A60 Microplus Nanoflow Cytometer (Apogee Flow Systems Inc.). Prior published work from our lab has already established technical settings to achieve data linearity using this system.^51,67^ Size gating was determined by calibration beads to observe smaller (180-300 nm) and larger particles (300-1000 nm), as previously published.^67^ Instrument settings included sheath pressure at 200 mbar, flow rate of 1.50 µL/min for 130 µL and lasers at 70 mW (405 nm), 70 mW (488 nm), and 70 mW (638 nm). Light scatter of events was produced using the 405 nm laser, with thresholds to eliminate background noise of 37 a.u. for small angle light scatter (SALS) and 40 a.u. for large angle light scatter (LALS). Photomultiplier tube (PMT) voltages were set at 260 V at LALS, 340 V at SALS, 525 V at L405-Blu, 525 V at L488-Grn, and 575 V at L638-Red. For each nanoflow experiment, samples were run in triplicate from 3 replicates.

### Statistical analysis

Statistical analysis was conducted using GraphPad Prism 10 (GraphPad Software, USA). All data is represented as mean ± SEM. Unpaired t-tests and one-way or two-way ANOVAs with Tukey’s multiple comparisons test were used for statistical analysis. Statistical significance was defined as \**p* < 0.05, \*\**p* < 0.01, or \*\*\**p*<0.001.

