## Supplemental Figures for "Amyloid beta is released by lysosomal exocytosis from hiPSC-derived neurons"

*Generation of iNeurons from hiPSCs*

To verify the neural yield of matured neurons, immunofluorescence staining was performed for nestin, a neural progenitor cell marker, and MAP2, a neuronal cell maker. Prior to differentiation and maturation, 92.82% of wild-type cells were nestin positive, whereas only 34.50% of wild-type cells were MAP2 positive ($p<0.001$). Likewise, 80.77% of APP-Swe cells stained positively for nestin while only 25.74% of APP-Swe cells stained positively for MAP2 ($p<0.001$). After 7 days of differentiation and 21 days of maturation, 67.13% of wild-type cells were MAP2 positive and 23.23% were nestin positive ($p=0.006$). Similar findings were detected among neural differentiated APP-Swe cells, where 68.2% of cells were MAP2 positive and 23.30% were nestin positive ($p=0.005$). In undifferentiated wild-type and APP-Swe cells, the cell bodies appeared larger with fewer distinct processes. Upon differentiation, MAP2 staining revealed elongated cells were with extended processes emanating from smaller, more triangular cell bodies.

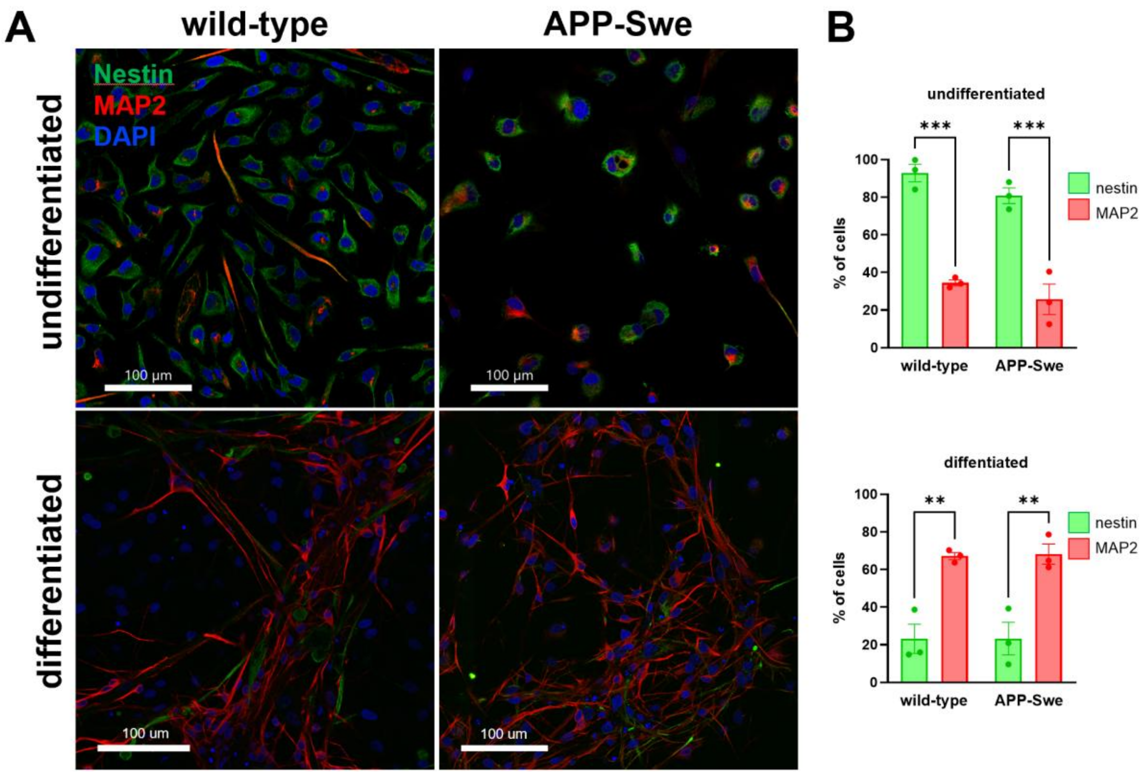


**Supplementary Figure 1 Immunofluorescence staining of nestin and MAP2 in wild-type and APP-Swe iNeurons**

A) Representative images of cells stained for nestin (green) and MAP2 (red) before and after 7 days of differentiation and 21 days of maturation. Nuclei were counterstained with DAPI (blue). B) Statistical analysis of the relative % nestin and MAP2 positive cells was performed using the filament tracer function on Imaris. All results are presented as mean ± SEM. Asterisks indicate significant changes according to one-way ANOVA and post hoc Tukey test. $^{**}p<0.01$ $^{***}p<0.001$

<u>*Silencing of Rab27b and munc13-4*</u>

In wild-type cells, relative steady-state protein levels of Rab27b decreased to an average of 0.0603 (p=0.002) compared to shScramble control following silencing. Relative steady-state protein levels of munc13-4 also decreased to an average of 0.0925 (p<0.001) compared to shScramble control following silencing in wild-type iNeurons. Similarly, in APP-Swe cells, relative steady-state protein levels of Rab27b decreased to an average of 0.0689 (p<0.001) compared to shScramble control following silencing. Relative steady-state protein levels of munc13-4 also decreased to an average of 0.135 (p<0.001) compared to shScramble control following silencing in APP-Swe iNeurons.

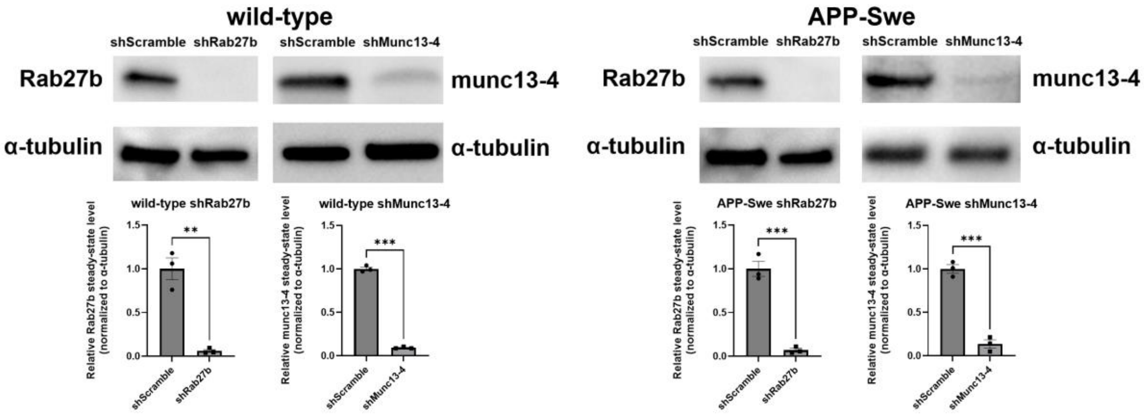


**Supplementary Figure 2 Silencing of Rab27b and munc13-4 in wild-type and APP-Swe iNeurons**

Western blotting with quantification of relative steady-state protein levels to verify silencing of Rab27b and shMunc13-4. All results are presented as mean ± SEM. Asterisks indicate significant changes according to unpaired t-test. **p<0.01 ***p<0.001
